# Exposing the Molecular Reaction Blind Spots of LLMs with PathwayQA

**DOI:** 10.1101/2025.08.12.669911

**Authors:** Gowri Nayar, Kristy A. Carpenter, Delaney A. Smith, Betty Xiong, Russ B. Altman

## Abstract

Proteins mediate a large portion of cellular activity, and understanding protein pathways can yield novel biological insights. Large language models (LLMs) have become increasingly adept at performing inference tasks across different fields of science and engineering. These models could facilitate the analysis of protein networks and help generate hypotheses about protein interactions in a scalable and accessible manner. However, the performance of LLMs in inferring protein-mediated biochemical reactions remains understudied. Here, we evaluate nine LLMs in reasoning over protein pathways included in the curated Reactome database. We find that all nine models struggle to infer products of a reaction when given reactants and enzymes. GPT-4o mini performed the best with a median recovery score of 0.6667, but no model surpassed the baseline strategy of parroting reactants back as predicted products. Most LLMs also performed poorly when inferring whether a protein pathway is associated with a human disease, with an average accuracy of 0.5980. DeepSeek 7B Chat performed the best with an accuracy of 0.9100. This study highlights an area where LLMs still struggle to make correct inferences and provides an opportunity for further work in developing biological LLMs. We also provide a novel question-answer dataset, PathwayQA, which is based on the Reactome database. PathwayQA can be used to benchmark and improve model performance on reasoning over protein-interaction networks. PathwayQA is available at https://github.com/Helix-Research-Lab/PathwayQA.

## 1. Introduction

Protein pathways organize biochemical reactions into coherent processes that underlie cellular function, with reactions as their granular components. These pathways are key for identifying disease mechanisms and potential drug therapeutic targets.^1^ Pathways have been characterized via biological experimentation and catalogued in databases over decades. Databases, such as the Kyoto Encyclopedia of Genes and Genomes (KEGG),^2^ the Search Tool for the Retrieval of Interacting Genes/Proteins (STRING),^3^ and Reactome,^4^ have been curated to collate protein pathways, which are composed of sequential chemical reactions. Reactome is one of the largest and most comprehensive manually curated databases of human protein pathways, providing high confidence in the stored reactions. Pathways are represented as hierarchical, sequential networks that vary in complexity;^4^ well-studied pathways (e.g., glycolysis) have abundant literature support, while others are created from lower-confidence evidence. Because these resources are incomplete and biased toward well-characterized entities, inferring novel reaction and pathway relationships remains challenging, limiting functional annotation across the full human proteome.^5^ Furthermore, accessing the data at a full-proteome scale requires technical knowledge to query and structure effectively, reducing accessibility for many biologists. Therefore, scalable computational methods are necessary to infer new functional relationships.

Large language models (LLMs), built on the transformer architecture, efficiently process sequential data via self-attention and can often perform well as few-shot learners without extensive fine-tuning.^6^ These capabilities have enabled their success across many downstream tasks in the biomedical domain. Recent advances have improved their performance on complex problems such as summarizing published articles,^7^ biomedical question answering,^8^ protein design,^9^ and passing multiple-choice medical board exams.^10^ Yet their capacity to reason over complex, multi-step biological systems—like protein pathways involving promiscuous entities and sequential molecular transformations—varies.^11^

Prior work has begun to tackle the problem of LLM reasoning over protein interactions. ProLLM^12^ has made advances by introducing a Protein Chain of Thought framework, where it transforms protein-protein interaction data into natural language prompts. BioMaze^13^ has introduced a dataset of 5.1k pathway problems and PathSeeker, an LLM agent that reasons through interactive subgraphs of biological pathways. However, this prior work does not model the chemical transformations that occur during protein interactions. Specifically, these papers do not address the problem of predicting the products of reactions or pathway disease associations, as documented in Reactome.^4^ These capabilities are fundamental for contextualizing novel findings and generating reasonable hypotheses based on biochemical understanding and disease context.

To fill this gap, we built PathwayQA, a benchmark question and answer (QA) dataset based on Reactome. PathwayQA consists of two subtasks: predicting reaction outputs and linking pathways to diseases. The datasets are packaged as easy-to-use prompt-answer pairs and used to evaluate both biologically tuned open-weight LLMs (BioGPT^14^ and BioMedLM^8^) and general-purpose models (Claude 3.5 Haiku,^15,16^ GPT-4o mini,^17^ DeepSeek 7B Chat,^18^ Gemma 7B Instruct,^19^ Llama3.1 8B Instruct,^20^ Mistral 7B Instruct,^21^ and Qwen1.5 7B Chat^22^). GPT-4o mini achieves the highest recall on reaction output generation, DeepSeek the highest accuracy on disease association, and Mistral the best accuracy for disease name generation. Despite that, all nine models perform poorly overall: the top average recall for reaction outputs is 0.6667, and the top accuracy for disease name generation is < 0.6. Therefore, this work underscores the need to fine-tune models using the proposed benchmarks. PathwayQA is available at https://github.com/Helix-Research-Lab/PathwayQA.

## 2. Methods

We systematically curate Reactome into a question-answer dataset, and evaluate current LLMs for their ability to generate data related to protein pathways (Figure 1).

**Fig. 1.**
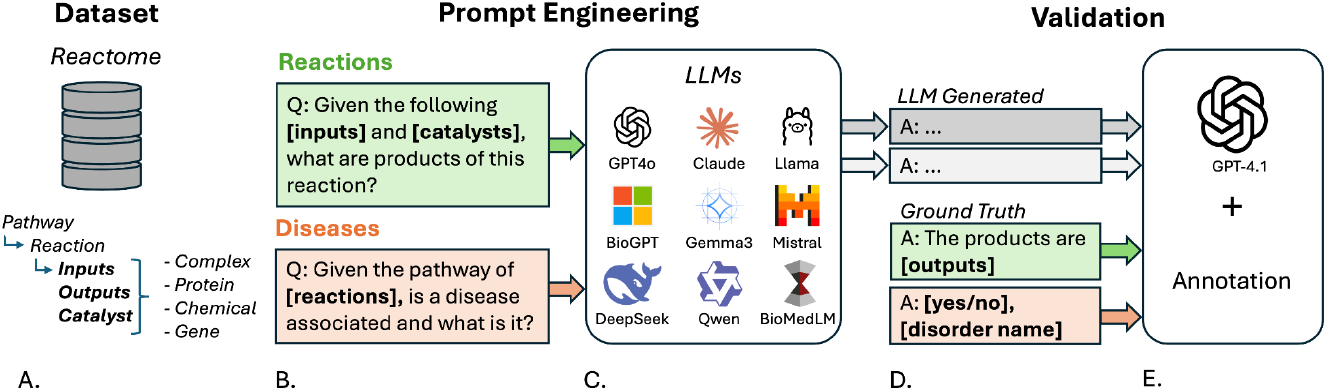
Pipeline for evaluating LLMs on protein-pathway reaction and disease prediction. **(A)** We extract reactions, pathways, and disease associations from Reactome to **(B)** build two QA benchmarks (reaction outputs and pathway-disease links). **(C)** We prompt nine LLMs (general purpose and fine-tuned) on both datasets, and **(D)** validate their answers against Reactome ground truth using GPT4.1, with comparison to human annotations.

### 2.1. Dataset curation from Reactome

We compiled a set of 2,306 pathways, which consist of 15,168 reactions, from Reactome. Each reaction involves reactants, enzymes, and products, each of which can be proteins, organic small molecules, ions, gene products, or a complex of multiple entities together.

We used the Reactome API to generate a map linking each Reactome entity to its corresponding reference-database ID and name. For proteins, the reference database is UniProt,^23^ for chemical compounds it is ChEBI,^24^ and for gene products it is Ensembl.^25^ We decomposed each complex into its sub-entities and stored the mapping between the Reactome entity ID for the complex and the corresponding reference identifier for each sub-entity.

We applied this mapping to annotate every reactant, product, and enzyme of each reaction with its reference-database identifier and name. We consolidated all reactions into a structured JSON file. For each reaction, it includes the Reactome reaction ID, the name of the reaction, and the reactant, product, and enzyme entities (each with Reactome IDs, reference database IDs, and names). We maintained the Reactome ID for each element so that future users may use it to query the API for necessary metadata.

We also retrieved Reactome’s pathways, which consist of an ordered sequence of reactions, along with the pathways’ disease annotations. For each pathway, we generated a structured JSON file that denotes the Reactome pathway ID, name of pathway, disease association, pathway context (biological relevance of the pathway), disease name (if applicable), and an ordered list of its reaction components.

We construct the two benchmark question-answer (QA) datasets that make up PathwayQA using the two files. We used the reaction file to create the reaction subtask of PathwayQA, where the reactants and enzymes are given in the prompt and the LLM is asked to generate the products. We used the pathway file to create the disease subtask of PathwayQA, where the pathway and pathway context are in the prompt, and the LLM is asked to generate the disease association and disease name, if applicable.

### 2.2. Prompt engineering to create the QA dataset

We performed prompt engineering to iteratively develop prompts for both tasks. For the reaction output generation task prompts, we first suggested to the model that it is an expert in biochemistry, explained the structure for providing outputs, and gave two examples of completed reactions (reactants, enzymes, products). For each reaction, we generated a query that gave the reactant and enzyme entities in the prompt and asked the LLM to generate the outputs. The answers associated with these prompts consisted of the true set of product entities, with both names and reference identifiers. For the disease association task, we generated a prompt that provided the pathway identifier and pathway context extracted from Reactome and asked the LLM to infer whether the pathway is associated with a disease, and if so, which disease. The answers associated with these prompts were a yes/no label of whether the pathway is known to be associated with a disease, followed by the disease name if available. We also tested prompts with one example (“one-shot”) and no examples (“zero-shot”) for the reaction product generation task. For disease association, we only tested one-shot prompts due to the simpler structure of the task. Example question/answer pairs for both tasks are present in Supplementary Text S1-2.

### 2.3. Evaluated Models

We evaluated recent state-of-the-art LLMs of comparable size, including general-purpose models trained on broad text corpora, and biomedical models, which are smaller and updated less often. We prioritized small, freely available models due to their accessibility for research and included two affordable paywalled models to compare performance.

The nine LLMs we evaluated were: (1) BioGPT,^14^ a model pretrained and fine-tuned on biomedical literature for domain tasks and the smallest model evaluated; (2) BioMedLM,^8^ a GPT-style model pretrained on biomedical text for question answering; (3) Claude 3.5 Haiku,^15,16^ a paid general-purpose model that is the cheaper and lighter version of the popular Claude Sonnet; (4) DeepSeek 7B Chat,^18^ a general-purpose, decoder-only model optimized for speed and resource use; (5) Gemma 7B Instruct,^19^ a general-purpose, decoder-only model that is the lighter version of the popular Google Gemini; (6) GPT-4o mini,^17^ a paid general-purpose model that is the cheaper and lighter version of OpenAI’s popular and versatile GPT-4o; (7) Llama3.1 8B Instruct,^20^ a general-purpose open source model finetuned for instruction; (8) Mistral 7B Instruct,^21^ a general-purpose open source model tuned on instruction data (prompt-response, summarization, and QA); (9) Qwen1.5 7B Chat,^22^ a general-purpose decoder-only model finetuned for instruction and conversational safety. See Supplementary Table S3 for architectural detail for each model and Supplementary Text S4 for details on run-time parameters.

### 2.4. Evaluation

#### 2.4.1. Evaluate reaction output generation with GPT 4.1 and human annotation

To perform LLM-as-a-judge,^26^ we used a larger, newer, and performance-optimized LLM, OpenAI’s GPT4.1,^27^ to determine if the outputs of the 9 selected models matched the true reaction products. We prompt GPT4.1 to mark each ground-truth entity (protein, chemical compound, gene product, complex) as True if it appears in the generated output, and False otherwise. We then compute a ‘reaction score’ as the fraction of ground-truth entities recovered. ‘Pathway scores’ are the average of their constituent reaction scores.

To ensure that GPT4.1 evaluations were of satisfactory quality, we manually graded a subset of the generated responses against the answers extracted from Reactome. We selected 10 random queries from each model and had two biosciences PhD students independently evaluate the model-generated answers. We repeated this process twice for additional depth in coverage (total = 180 manually-annotated queries). For each query in our manually-annotated set, we determined the fraction of the products listed in the ground-truth answer that are present in the generated answer; a protein complex was counted as a single product entity. Each pair of human graders discussed their scores to resolve major discrepancies in grading style, although minor differences were preserved to reflect variance in rule interpretation (for a complete set of evaluation criteria, see Supplementary Text S5). After finalizing a consistent rubric, we computed the Spearman correlation between each pair of graders.

We created a validation set from 30% of the manually-annotated queries with which to engineer the GPT4.1 evaluation prompt. We iterated on the instruction prompt until GPT4.1 was able to replicate human grading on the validation set to a sufficient threshold of similarity (above r = 0.8 Spearman correlation).^28^ The final prompt is included in Supplementary Text S6. We ran the GPT4.1 grading query over the remainder of the manually-annotated queries and again calculated the Spearman correlation between the human annotations and GPT4.1 annotations. Our correlation met the threshold, so we used the GPT4.1 grader to evaluate the unannotated generated data.

We measure reaction output generation with recall: the fraction of ground-truth output entities in the generated output (0 = none, 1 = all). We compared model performance to a baseline strategy of parroting the reactants of the query back as the predicted products. To assess extraneous content, we compare the length of each generated answer to the ground-truth answer by computing the Spearman correlation between their word counts; a higher correlation indicates the model’s output length better matches the true answer and thus contains fewer extraneous additions.

#### 2.4.2. Evaluate the impact of proteins’ promiscuity on model scores

A promiscuous protein can react with different molecules and thus participate in different reactions. The number of times the protein appears in the full Reactome dataset, across all reactions, can be a metric for promiscuity. More promiscuous proteins are found in more reactions and thus are more frequently in the literature and the training data. This can impact an LLM’s performance, as it has seen more instances of the promiscuous proteins in training^29^. To measure the level of promiscuity for each reaction and pathway, we find the number of times each component protein is present in the dataset and normalized by the number of proteins in the reaction or pathway, which we define as the average protein occurrence.

To assess the impact of promiscuous and well-studied proteins, we correlate model scores with the average protein occurrence of a reaction’s and pathways’ entities in the full Reactome dataset. For each reaction and pathway, we compute the mean occurrence of its entities, normalized by the total number of entities, to measure how common they are. We then analyze how this average correlates to model performance.

#### 2.4.3. Evaluate disease association generation with GPT 4.1 and regular expressions

To evaluate disease-association queries, we first used regular expressions to extract “Yes/No” answers from LLM outputs and compared them to Reactome labels. If they matched, the result was marked ‘True’. We then had GPT-4.1 perform the same task and compared its scoring using accuracy, F1, and MCC. Given high agreement between methods (accuracy > 0.95), we used GPT-4.1 for final evaluation, as its reasoning better captured edge cases missed by string matching.

For disease name generation, we evaluated only disease-associated pathways. We used approximate string matching—marking outputs ‘True’ if the correct disease name appeared in the response—and also asked GPT-4.1 to judge whether the generated name was a fuzzy match to the Reactome label. GPT-4.1 was used to detect biologically consistent but textually varied responses (e.g., “cancer” vs. “multiple myeloma”). Again, high agreement with pattern matching (accuracy > 0.95) led us to adopt GPT-4.1 scoring for final evaluation.

## 3. Results

### 3.1. Generated answers from general-purpose models are more accurate than fine-tuned models

We prompted all 9 models with the queries from both datasets and generated the set of answers, reaction outputs, and disease annotations. In Table 1, we show the generated outputs for two arbitrarily selected reactions for one general-purpose model, GPT-4o mini, and one fine-tuned model, BioGPT. The general-purpose model gets both the name of the product as well as the reference ID correct, while the fine-tuned model only gets the name correct. For the first reaction, both models score a 1 (all ground-truth entities are present in the generated output); GPT-4o mini correctly identifies the name, subcellular localization, and the reference ID, while BioGPT only gets the name correct. In the second example, GPT-4o mini scores a 1 and BioGPT scores a 0 (Table 1).

**Table 1.** Sample of generated reaction outputs for two randomly selected reactions for GPT-4o mini (highest-performing general-purpose model) and BioGPT (highest-performing fine-tuned model). Correct parts of generated outputs are colored in teal, and incorrect portions are colored in red.

| Ground-Truth | GPT 4o-mini | BioGPT |
| --- | --- | --- |
| *Catalyst:* botG LC:Zn2+ [cytosol] (complex of BoNT/G LC [cytosol] (UniProt: Q60393), Zn2+ [cytosol] (ChEBI: 29105))<br>*Reactants:* VAMP1 [synaptic vesicle membrane] (UniProt: P23763)<br>**Products:** VAMP1(1-83) [cytosol] (UniProt: P23763) | Cleaved VAMP1 [cytosol] (UniProt: P23763) | ADP-ribosylation of VAMP (uniProts: ChEB: 20861, ChEPP: 260697). |
| *Catalyst:* N/A<br>*Reactants:* p-T,4S-IRF3:p-T,4S-IRF3 [nucleoplasm] (complex of p-4S,T404-IRF3 [nucleoplasm] (UniProt: Q14653)), CREBBP,EP300 [nucleoplasm] (complex of CREBBP [nucleoplasm] (UniProt: Q92793), EP300 [nucleoplasm] (UniProt: Q09472))<br>**Products:** p-T,4S-IRF3:p-T,4S-IRF3:CREBBP, EP300 [nucleoplasm] (complex of p-4S,T404-IRF3 [nucleoplasm] (UniProt: Q14653), CREBBP [nucleoplasm] (UniProt: Q92793), EP300 [nucleoplasm] (UniProt: Q09472)) | p-T,4S-IRF3:CREBBP,EP300 [nucleoplasm] (complex) | ADP, GDP, GTP, GMP, IMP, UDP, CDP, UMP, UTP, CTP, dCTP |

Supplementary Figure S7 shows that the general-purpose models more consistently score higher than the fine-tuned models, across easy and hard reactions and pathways. Fine-tuned models only succeed on ‘easy’ queries, where all models were able to produce the correct answer.

DeepSeek and BioGPT, the general-purpose and fine-tuned models, respectively, had the best performance for disease name generation. In both cases, DeepSeek correctly classified the pathway while BioGPT was incorrect (Table 2). DeepSeek also correctly generated the disease name in the first example. Although it was incorrect in the second example, it still represents a related phenotype, as DeepSeek generated “antimicrobial resistance” when the ground truth disease is a bacterial infection.

**Table 2.** Sample generated reaction outputs for two randomly selected pathways (with disease annotation) for DeepSeek (highest-performing general-purpose model) and BioGPT (highest-performing fine-tuned model). Correct parts of generated outputs are colored in teal, and incorrect portions are colored in red.

| Ground-Truth | DeepSeek | BioGPT |
| --- | --- | --- |
| *Pathway context:* 2-LTR circle formation<br>**Answer: yes, Human immunodeficiency virus infectious disease** | yes, Human immunodeficiency virus infectious disease | No |
| *Pathway context:* Action of antimicrobials and antimicrobial resistance<br>**Answer: yes, bacterial infectious disease** | yes, Antimicrobial resistance | Yes, Followed by the same name. |

### 3.2. GPT4.1 grading is scalable and consistent with human grading

The Spearman correlation coefficient between each pair of human raters on the two manually-annotated sets of 90 queries was 0.9116 (*p* = 4.97e-36) and 0.9475 (*p* = 1.22e-45), respectively. The Spearman correlation coefficient between human scores and GPT4.1 scores was 0.8069 (*p* = 8.67e-14) on the validation set (n = 54 queries) and 0.8362 (*p* = 1.28e-48) on the full manually-annotated set (n = 180 queries).

For disease association, the accuracy of GPT4.1 compared with text-based pattern matching was 0.9584 across all models, with an F1 score of 0.9661 and an MCC of 0.9149. The only model that had an accuracy < 0.95 was BioMedLM (accuracy = 0.67887, F1 = 0.4798, MCC = 0.4798). There were 316 pathways associated with disease (positive label), while 1,990 pathways had no disease association (negative label). For disease names, the agreement between text-based pattern matching and GPT4.1 was 0.9866 across all models, with an F1 score of 0.9892 and an MCC of 0.9791. Of the 316 disease-associated pathways, 312 pathways had a disease name listed in Reactome.

### 3.3. Models did not outperform the baseline strategy for identifying output generations

The distributions of recall scores varied across models, with Claude and GPT-4o mini performing the best in the two-shot and one-shot settings, and Llama and Mistral joining GPT-4o mini as the top performers in the zero-shot setting (Figure 2). BioGPT, BioMedLM, DeepSeek, and Qwen had median recall scores of 0 in all three prompt settings. While the difference in performance between the two-shot and one-shot setting was small and of variable directionality, the zero-shot setting often had slightly improved performance. No model achieved excellent performance on this task, with the baseline strategy of parroting the query reactants back as predicted products outperforming all methods. The highest median recall score was 0.6667 (tie between GPT-4o mini zero-shot, Llama zero-shot, and Mistral zero-shot), and the interquartile ranges of the best-performing models averaged around 0.75 (Figure 2).

**Fig. 2.**
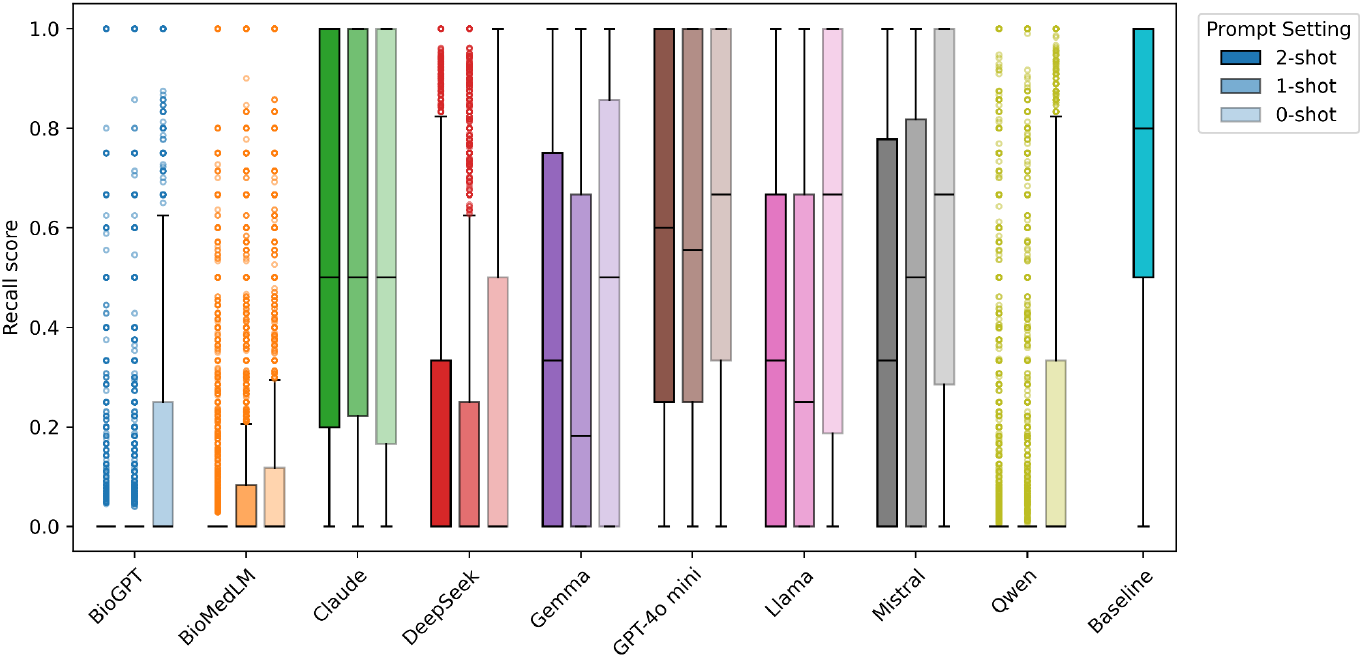
LLM performance evaluation for reaction products recall. Each model is denoted by a different color and grouping of boxes; within each grouping, from left to right (and with decreasing opacity), are the results for the two-shot, one-shot, and zero-shot settings. The teal box on the far right, labeled “Baseline,” is the performance when the reactants of the query reaction are parroted back as the predicted products. Each box spans the first quartile to the third quartile, with whiskers extending to 1.5x the interquartile range from the box. Outliers beyond the whiskers are shown as individual dots. The median of each distribution is shown as a black line.

### 3.4. Model score decreases when the reaction includes more promiscuous proteins

Model recall scores decrease with increased protein promiscuity (Figure 3). A more promiscuous protein (interacting with different molecules in different reactions) will occur more frequently in the Reactome dataset; thus, reactions and pathways with promiscuous proteins will have a higher average protein occurrence. Across all models, we see that as the average protein occurrence per reaction increases, the models’ recall scores decrease. Per reaction, the models with higher overall recall scores (Claude and GPT-4o mini) exhibit a more significant decrease in recall score with an increase in average protein occurrence. In contrast, models with lower overall recall scores (BioGPT and BioMedLM) showed less variance as the average protein occurrence increased. Supplement Figure S9 shows the comparison between the model recall score and the average protein occurrence per pathway. The decreasing trend was not reflected in the pathway recall scores; across all models, the scores remained relatively constant as the average protein occurrence per pathway increased.

**Fig. 3.**
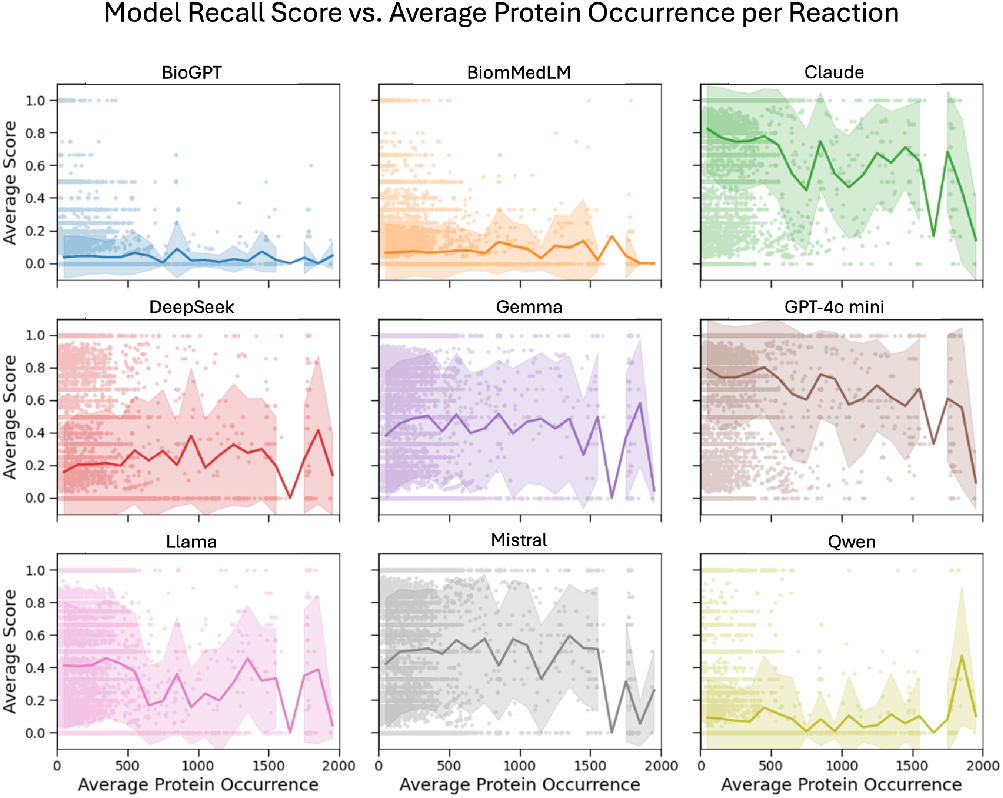
Effect of protein occurrence on model performance for reaction recall scores. Reaction-level recall scores are plotted against the average occurrence of the reaction’s proteins in the full Reactome dataset. Across all models, scores decline as protein occurrence increases, suggesting that frequently occurring (promiscuous) proteins introduce ambiguity by participating in multiple reactions.

### 3.5. Models generate a consistent amount of text irrespective of the true answer length

Most models exhibited weak or no correlation between true answer length and generated answer length (Spearman’s r ranging from 0.00 for Gemma to 0.28 for GPT-4o mini), showing that most models have a standard length of output text regardless of the ground-truth output length (Figure 4). BioGPT stands out with a high correlation of r = 0.71. BioGPT, BioMedLM, Claude, DeepSeek, and Mistral tend to output more text than required to answer correctly (Figure 4).

**Fig. 4.**
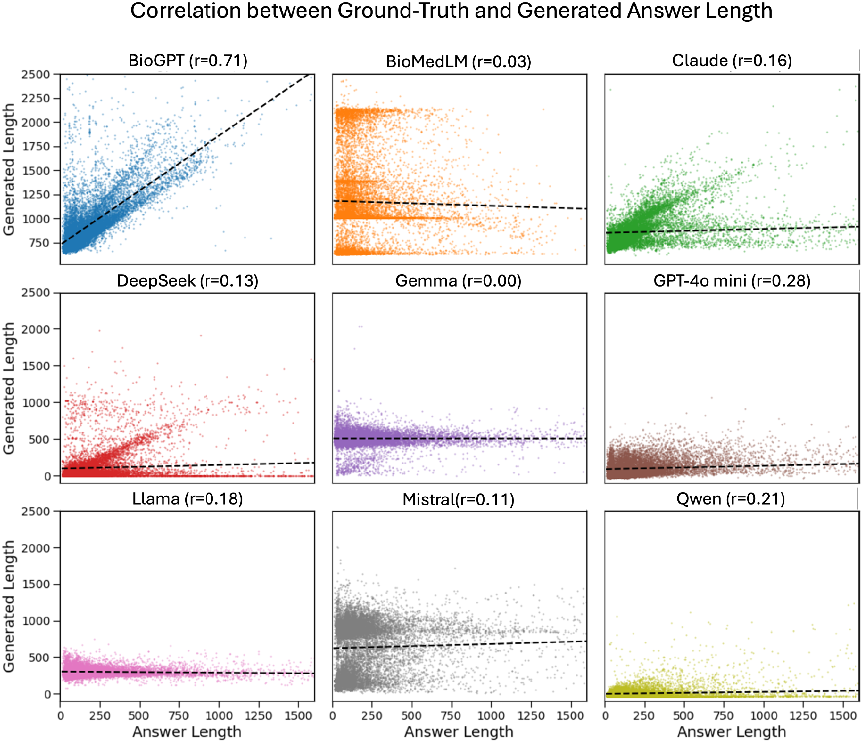
Correlation between generated and ground-truth output lengths for reaction prediction. For each model, we plot the number of words in the generated output versus the number of words in the ground-truth output for all reaction prompts. Dashed lines indicate linear regression fits, and the Spearman correlation coefficient (r) is reported in the title of each subplot. BioGPT shows the strongest correlation (r = 0.71). The other models show weak or no correlation between ground-truth outputs and generated outputs.

### 3.6. Models vary in their ability to infer disease-pathway associations and disease names

The average accuracy across all models for disease-pathway association was 0.5980 with an F1 score of 0.3230 and an MCC of 0.1972. The best performing model, by all metrics, was DeepSeek (accuracy (acc) = 0.9100, F1 = 0.6391, MCC = 0.5912) (Figure 5A). Mistral (acc = 0.8184, F1 = 0.5333, MCC = 0.4661) and GPT-4o mini (acc = 0.7990, F1 = 0.545, MCC = 0.499) also had an F1 score above 0.5. BioGPT had the worst F1 score and MCC (acc = 0.802, F1 = 0.258, MCC = -0.0720).

**Fig. 5.**
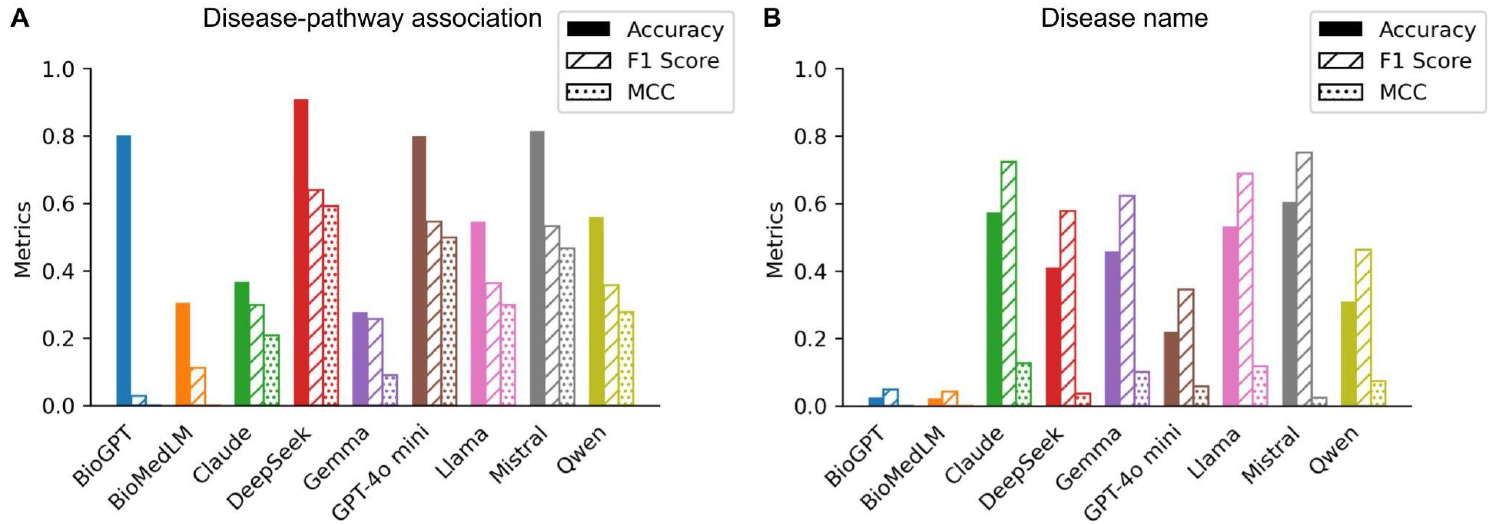
LLM performance evaluation for inferring **A:** disease-pathway associations and **B:** disease names. Specifically, panel A shows model performance in inferring whether or not a protein pathway in Reactome is associated with a disease. In panel B, the models are asked to generate the name of the associated disease, and a binary score is assigned based on whether the disease names are consistent at the biological level across Reactome and the LLM-generated answer. Accuracy is shown as a solid bar, the F1 score as a hatched bar, and the MCC as a dotted bar.

For disease name evaluation, the average accuracy across all models was 0.3512 with an F1 score of 0.5310 and an MCC of 0.0101. The best performing model was Mistral with 0.6041 accuracy, an F1 score of 0.7514, and an MCC of 0.024 (Figure 5B). Claude (accuracy = 0.5730, F1 = 0.724, MCC = 0.1284) and Llama (accuracy = 0.532, F1 = 0.623, MCC = 0.1180) also had an accuracy above 0.50. BioGPT (accuracy = 0.0254, F1 = 0.0491, MCC = -0.570) and BioMedLM (accuracy = 0.022, F1 = 0.043, MCC = -0.0596) had accuracies < 0.05 (Figure 5B).

## 4. Discussion

As LLMs grow in size and capability, so does the potential for these tools to accelerate biological research. To assess whether their outputs are sufficiently accurate and reliable, domain-specific and tractable benchmarks are essential. We address this need with three main contributions: (1) we introduce PathwayQA, a benchmark textual, question-answer dataset derived from the Reactome database, (2) we evaluate nine LLMs (general purpose and fine-tuned) on their ability to generate correct reaction outputs and disease associations, and (3) we demonstrate the need for improved models specifically for reaction and pathway generation.

In our evaluation, the largest and most expensive models performed the best in the one- and two-shot settings for the reaction product generation task, and the smallest and oldest models performed the worst in all prompt settings. This is likely due to the increase in reasoning capability with more parameters. The disparity is further exacerbated by the potential orders of magnitude difference in parameter count between BioGPT/BioMedLM and GPT-4o mini/Claude, a discrepancy that OpenAI and Anthropic obscure. Gemma, Llama, and Mistral outperformed DeepSeek and Qwen, likely due to their tendency to generate extra reactions. In contrast, DeepSeek and Qwen tended to have brief responses that matched the expected output format. We demonstrated that regurgitating the reactants outperformed all models, suggesting significant room for improvement in this task, as models that performed well likely capitalized on the similarity between products and reactants. Relatedly, the unexpected increase in performance with zero-shot prompting may stem from the model more easily repeating reactants instead of copying unrelated examples from the prompt (as is common in the outputs from the two-shot and one-shot prompts). Parroting back reactants likely did well because in some cases, an entity is both a product and a reactant, and in some cases, the subtle changes between the reactant form and the product form were not caught in our grading scheme. We kept our grading scheme lenient because of the amount of hallucination and poor overall performance of the assessed models.

The top-performing model in identifying disease associations was DeepSeek, followed by GPT-4o mini and Mistral. While GPT-4o mini also performed well in the product generation tasks, DeepSeek only performed well in inferring disease associations (accuracy above 0.9). This version of DeepSeek was tuned for reasoning ability, which may explain its enhanced performance.^18^ With disease name generation, no model had an accuracy above 0.8. Due to prompt length constraints, we only provided the pathway ID and pathway context in the query, as opposed to the chain of all proteins in the pathway. It is possible that this did not give the model sufficient context to identify the pathway.^30^

However, evaluating model performance on accuracy alone is not sufficient for handling the class imbalance in the disease-pathway association and disease-name tasks. Most pathways (86.3%) have no associated disease, creating a strong bias toward negative labels. As a result, metrics like accuracy and F1 score could overestimate performance. To address this, we used a combination of metrics—including the Matthews Correlation Coefficient (MCC)—to assess each model’s ability to handle imbalanced data more accurately. Only DeepSeek achieved a strong correlation (MCC > 0.8) for disease association. In the disease-name task, nearly all disease-associated pathways (except 4) had a specific name. For an LLM’s output to be marked correct, it needed to generate a functionally equivalent disease name—a non-trivial task unlikely to be answered correctly by chance. Therefore, in this case, accuracy served as a more meaningful metric than class-sensitive alternatives.

When evaluating model performance in product generation via manual annotation, we observed that each model had its own idiosyncrasies. In the product prediction task, Claude and GPT-4o mini frequently repeated the enzymes and reactants of the queried reaction, which at times led to an artificially high recall score due to reactants and products being similar. BioMedLM attempted to force the output to conform to text styles found in published articles. The hallucinations produced by BioMedLM were so extensive and nonsensical that GPT4.1 struggled to evaluate the results in the disease association task. For a human reader, the hallucination and regurgitation produced by these models frequently obscured the correct answers in the generated text. Sample outputs of these texts are available in Supplementary Text S8.

Most of the models maintained a strict format for the generated text and constant length of output (Figure 4). Claude, GPT-4o mini, Llama, Mistral, and Gemma tended to add extra reactions in the same pattern at the end of the generated output, decreasing their correlation with ground-truth output length. BioGPT, DeepSeek, and Qwen did not add extraneous text as frequently, and their output lengths had a higher correlation to the ground-truth answer. Despite this, these three models performed poorly overall in generating correct reaction outputs (Figure 2), suggesting that they had likely learned to approximate output size based on superficial features—such as the number of input entities—rather than reasoning over the chemical transformation. In contrast, models such as GPT-4o mini showed more modest correlations but achieved higher accuracy, indicating better alignment with actual content rather than just output length.

Despite consistent formatting, LLMs showed inconsistent performance in capturing biochemical details. Reactions with a higher average protein occurrence (containing promiscuous proteins) had lower recall scores, likely due to the broader range of possible outputs. In contrast, reactions with rare proteins (lower occurrence rates in the dataset) scored higher, as there were fewer instances in the training data to choose from. The average protein occurrence rates per pathway did not affect the pathway recall score significantly, likely because the pathway recall score is the average of its component reaction scores, and therefore robust to single outliers. Furthermore, the models often identified common cofactors like ADP and GTP, likely due to repetition, pattern matching, or their frequent occurrence in reactions. However, they rarely recognized lone phosphates (Pi) as products. We found that the average protein occurrence (used as a measure of the proteins’ promiscuity) per reaction was an important parameter for the models’ recall scores. Models also frequently missed complex components and correct chemical identifiers (e.g., ChEBI, UniProt).

Due to these idiosyncrasies and varied outputs, human evaluation of model outputs was non-trivial, even when performed by domain experts. A key consideration was handling inconsistent biological nomenclature and representations: differences such as ubiquitinated versus non-ubiquitinated proteins, complex names versus their constituents, and varying naming conventions (e.g., 2-oxoglutarate vs. 2-OG, AdoHcy vs. S-adenosyl-L-homocysteine) complicated scoring and required domain expertise. Models also produced vague outputs like “peptide fragments” or “downstream signaling molecules,” making it unclear whether correct entities were inferred. Interrater agreement testing revealed that human graders varied in judgments, with the LLM judge (GPT4.1) sometimes showing greater consistency and stricter fairness, underscoring both the difficulty and potential of LLM-as-a-judge evaluation.

We also identify broader limitations in this study. We removed prompt repetition from model outputs but did not filter out extraneous reactions generated after the target query; a more robust evaluation would exclude these. Using an LLM as a judge adds uncertainty, as it may overlook errors a human expert would catch. While we thoroughly validate the LLM’s grading in this work, future improvements could include querying models multiple times to form consensus outputs and refining prompts further. The prompts for the initial QA tasks could also be refined further; all nine models were given the same prompts for consistency, but it is possible that some models would perform better with a modified prompt. Testing high-performance models at scale is computationally expensive, limiting our ability to fully assess their potential. Finally, the study relies solely on the Reactome database, which, while well-curated, is biased towards well-studied pathways and diseases.

Our results suggest that it may be useful to further tune LLMs on PathwayQA to potentially improve performance. Agentic LLMs – such as Biomni^31^ or Aviary^32^ – that are able to search the web and make API calls may be able to access Reactome directly and improve performance on these tasks. In any case, there is a clear need to further develop LLMs in the protein and chemical reaction space. We trust that our provided benchmark will assist in these activities.

## 5. Appendix

The code, data, and supplementary material accompanying this paper is available at github.com/Helix-Research-Lab/PathwayQA.

## Acknowledgments

GN is supported by NIH NLM F31LM014646; KAC is supported by NIH F31GM151783; DAS is supported by the Stanford Biochemistry Department; BX is supported by Australian-American Fulbright Future Scholarship; RBA is supported by NIH R35GM153195 and Chan Zuckerberg Biohub.

