## Supplement for "Exposing the Molecular Reaction Blind Spots of LLMs with PathwayQA": PSB LLM draft_supp.pdf

**Supplementary Material for "Exposing the Molecular Reaction Blind Spots of LLMs with PathwayQA"**

**Text S1.** Example question/answer pair for reaction product generation task in two-shot, one-shot, and zero-shot forms.

*Two-shot prompt*

You are an expert on chemistry and biology. Complete the following reactions with their products. Do not repeat the question, provide only the answer in a brief manner. Use the following example format

Enzyme: Hexokinase (UniProt: P19367)
Reactants: Glucose (ChEBI: 17234), ATP (ChEBI: 15422)
Products: Glucose-6-phosphate (ChEBI: 4170), ADP (ChEBI: 16761)

Enzyme: N/A
Reactants: GTP [mitochondrial matrix] (ChEBI: 37565), AMP [mitochondrial matrix] (ChEBI: 456215)
Products: GDP [mitochondrial matrix] (ChEBI: 58189), ADP [mitochondrial matrix] (ChEBI: 456216)

Enzyme: SFKs:p-KIT complex [plasma membrane] (complex of FYN [cytosol] (UniProt: P06241), LYN [cytosol] (UniProt: P07948), YES1 [cytosol] (UniProt: P07947), LCK [cytosol] (UniProt: P06239), p-7Y-KIT [plasma membrane] (UniProt: P10721))
Reactants: ATP [cytosol] (ChEBI: 30616), PI(3,4,5)P3 [plasma membrane] (ChEBI: 57836), VAV1 [cytosol] (UniProt: P15498)
Products:

*One-shot prompt*

You are an expert on chemistry and biology. Complete the following reactions with their products. Do not repeat the question, provide only the answer in a brief manner. Use the following example format

Enzyme: Hexokinase (UniProt: P19367)
Reactants: Glucose (ChEBI: 17234), ATP (ChEBI: 15422)
Products: Glucose-6-phosphate (ChEBI: 4170), ADP (ChEBI: 16761)

Enzyme: SFKs:p-KIT complex [plasma membrane] (complex of FYN [cytosol] (UniProt: P06241), LYN [cytosol] (UniProt: P07948), YES1 [cytosol] (UniProt: P07947), LCK [cytosol] (UniProt: P06239), p-7Y-KIT [plasma membrane] (UniProt: P10721))
Reactants: ATP [cytosol] (ChEBI: 30616), PI(3,4,5)P3 [plasma membrane] (ChEBI: 57836), VAV1 [cytosol] (UniProt: P15498)
Products:

*Zero-shot prompt*

You are an expert on chemistry and biology. Complete the following reactions with their products. Do not repeat the question, provide only the answer in a brief manner. Use the following example format

Enzyme: SFKs:p-KIT complex [plasma membrane] (complex of FYN [cytosol] (UniProt: P06241), LYN [cytosol] (UniProt: P07948), YES1 [cytosol] (UniProt: P07947), LCK [cytosol] (UniProt: P06239), p-7Y-KIT [plasma membrane] (UniProt: P10721))
Reactants: ATP [cytosol] (ChEBI: 30616), PI(3,4,5)P3 [plasma membrane] (ChEBI: 57836), VAV1 [cytosol] (UniProt: P15498)
Products:

*Answer*

ADP [cytosol] (ChEBI: 456216), p-VAV1:PIP3 [plasma membrane] (complex of PI(3,4,5)P3 [plasma membrane] (ChEBI: 57836), p-Y142,Y160,Y174-VAV1 [plasma membrane] (UniProt: P15498))

**Text S2.** Example question/answer pair for disease association task.

*Prompt*

You are an expert on chemistry and biology. Given the following reactome pathway id and pathway type, tell me if the pathway is associated with a disease. Do not repeat the question. Provide only the answer in the following format: yes/no, <disease_name>. If there is no disease association, write 'no, None'. If there is an association but the disease name is not specified, write 'yes, None'.
Pathway id: R-HSA-164843
Pathway context: 2-LTR circle formation
Answer: yes, Human immunodeficiency virus infectious disease

Pathway id: R-HSA-2978092
Pathway context: Abnormal conversion of 2-oxoglutarate to 2-hydroxyglutarate

*Answer*

Answer: yes, glioblastoma multiforme

**Table S3**. Large language models assessed in this study. *Note that neither Anthropic nor OpenAI have publicly released the parameter counts for Claude 3.5 Haiku or GPT-4o mini, respectively. Some sources estimate the parameter counts of both to be 8B, while others estimate both to be 175B.

| Model | Provider | Domain | Parameters | Access | Release year |
|---|---|---|---|---|---|
| BioGPT | Microsoft | Biomedicine | 347M | HuggingFace (Free) | 2022 |
| BioMedLM | Stanford/ MosaicML | Biomedicine | 2.7B | HuggingFace (Free) | 2022 |
| Claude 3.5 Haiku | Anthropic | General | ~8B* | Anthropic API (Paid) | 2024 |
| DeepSeek 7B Chat | DeepSeek | General | 7B | HuggingFace (Free) | 2023 |
| Gemma 7B Instruct | Google | General | 8.54B | HuggingFace (Free) | 2024 |
| GPT-4o mini | OpenAI | General | ~8B* | OpenAI API (Paid) | 2024 |
| Llama3.1 8B Instruct | Meta | General | 8.03B | HuggingFace (Free) | 2024 |
| Mistral 7B Instruct | Mistral AI | General | 7.24B | HuggingFace (Free) | 2023 |
| Qwen1.5 7B Chat | Alibaba | General | 7.72B | HuggingFace (Free) | 2024 |

**Text S4.** LLM usage details.

*BioGPT*
Inference framework: transformers package
Parameters: num_return_sequences=1, no_repeat_ngram_size=2, max_length = 1024
Notes: Some prompts were too long for the context window and were omitted.

*BioMedLM*
Inference framework: vllm package
Parameters: temperature = 0.0, top_p = 1.0, max_tokens = 512, stop = "###"
Notes: We omitted all prompts longer than 2000 characters as they were too long for the context window.

*Claude 3.5 Haiku*
Inference framework: Anthropic API
Parameters: temperature = 0.0, top_p = 1.0, max_tokens = 1024

*DeepSeek 7B Chat*
Inference framework: vllm package
Parameters: temperature = 0.0, top_p = 1.0, max_tokens = 512, stop = "\n"
Notes: In model instantiation, we had to downcast to dtype = "float16"; we omitted all prompts longer than 7000 characters as they were too long for the context window.

*Gemma 7B Instruct*
Inference framework: vllm package
Parameters: temperature = 0.0, top_p = 1.0, max_tokens = 128

*GPT-4o mini*
Inference framework: OpenAI API
Parameters: all defaults

*Llama3.1 8B Instruct*
Inference framework: vllm package
Parameters: temperature = 0.0, top_p = 1.0, max_tokens = 128

*Mistral 7B Instruct*
Inference framework: vllm package
Parameters: temperature = 0.0, top_p = 1.0, max_tokens = 512, stop = "</s>"

*Qwen1.5 7B Chat*
Inference framework: vllm package
Parameters: temperature = 0.0, top_p = 1.0, max_tokens = 512, stop = "\n"

**Text S5**. Reaction product manual grading evaluation criteria.

For each reaction, we first split the products at the component level. Complexes of several entities were considered a singular component. For manual annotation, we assigned 'partial credit' for the proportion of component entities named correctly by the LLM. For example, if one component of the products of a reaction was ADP, and the model listed ADP in its output, it would receive full credit (1/1). However, if the component was a complex (ex., GGC-RAB9:GTP) and the model only generated 'GTP' as part of its output, it would receive partial credit (1/2). These partial scores were then averaged over the reaction and subsequently over the pathway. We did not deduct credit for missing or incorrect chemical identifiers (ChEBI IDs or UniProt IDs). We did not deduct credit for incorrect subcellular localization. We considered the term correct, regardless of where it appeared in the generated text. We considered the gene, mRNA, and protein forms of the same entity equivalent. If a model generated an equivalent name for an entity that was not an exact match for the expected answer, it got full credit (e.g. 2-OG instead of 2-oxoglutarate). We were very lenient in giving credit even if there were differences in posttranslational modifications, though we enforced that whether a protein was ubiquitinated or not had to match the expected answer in order to receive credit.

**Text S6.** Final prompt for GPT4.1 grading of reaction product generation task.

You are given a reference answer (in the form of a list of products) and a generated answer. For each protein, chemical compound, gene product, or complex entity in the reference product list, determine whether the generated text answer contains each of the entities. Be lenient in determining the presence of the entity in the generated answer. Allow for the following discrepancies:

1. The subcellular localization, which is indicated in brackets (e.g. [periplasmic space], [cytosol], [nucleoplasm]), does not need to match
2. The exact text used to describe the name of the entities does not need to match perfectly; instead make sure the entities match in meaning between the generated and references answer. Small differences like phosphorylated residue index or partial charge do not matter. Be lenient.
3. Allow for differences between mrna, gene, or product forms of the entity. Consider all three version of the same entity as a match.
4. Allow for mismatches between the id number, if present. If the names in the reference list and generated text have the same meaning and the id number do not match, consider that entity as a match.
5. If the reference compound appears anywhere in the generated answer, it counts as a match, regardless of whether or not that section of the generation appears relevant.
6. When considering a complex, consider the listed constituents of the complex and not simply the name of the complex. A complex is denoted as '[complex name] [subcellular localization] (complex of [constituent], [constituent], ...)'. Do not consider the complex name and do not count the constituents of the complex separately from the complex. If only a small portion of the complex's constituents are present, do not say the complex is present.
7. If the generated answer is blank or NaN, then return False for all reference products.

Again, be very lenient, and allow for matches even if they are not exactly chemically or biologically alike.

Reference answer: {ref_output_list}
Generated answer: {generated}

Respond only with a valid JSON with the following structure (Do not include ANY explanatory text or code fences.):

{"Entity1": true, "Entity2": false, ...}

**Figure S7.** Comparison of model scores across reaction and pathway difficulty, defined by how many models predict them correctly (easy = many or all models correct; hard = few or none). Panel A: Average reaction-level score per model grouped by the number of models that got the reaction right. Reactions predicted correctly by all models have high scores across the board; for harder reactions (fewer models correct), GPT-4o-mini maintains relatively high scores while BioMedLM, BioGPT, and Qwen drop substantially. Panel B: Pathway-level averages, where a pathway is considered correct for a model if the mean score of its component reactions exceeds 0.5. The same pattern holds: GPT-4o mini sustains higher scores on difficult pathways, whereas BioMedLM, BioGPT, and Qwen underperform. No pathway is correctly predicted by all nine models.

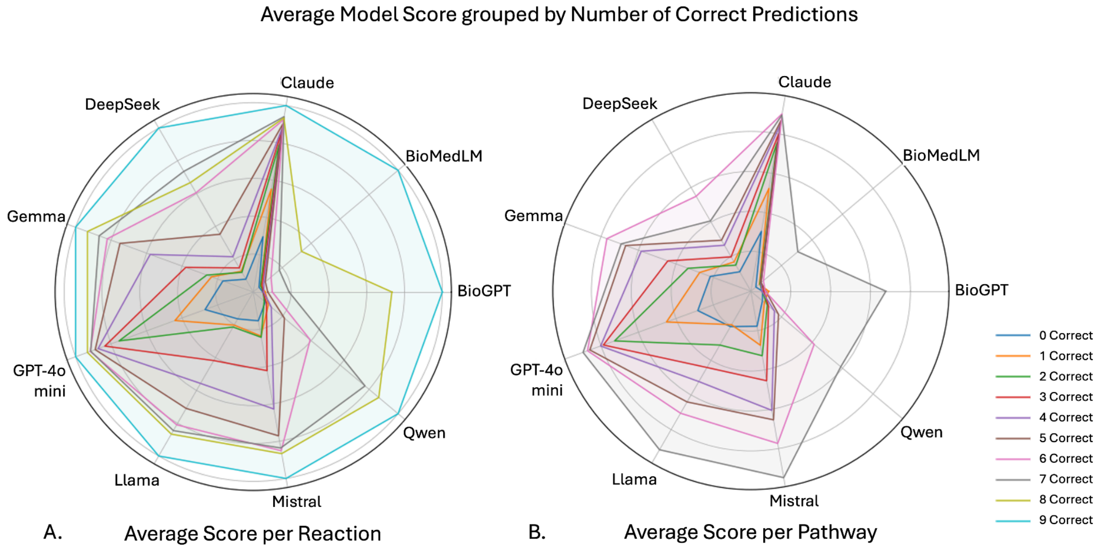


**Text S8.** Examples of reaction outputs/disease association generations with severe hallucinations and nonsensical output.

*Reaction product task, Example 1 (BioMedLM)*
\[ChEBI: 456216\] and p-PlexinA:P75601\] (*P75601\] (Figure 1, *P75601\] (*P75601\]

**Abbreviations:**

: **GDP, \[PMID: 28853896, **GTP, **R**as **g**lucy (UniProt: P2RY1 [**](#fn1){ref-type="fn"}, **t**ris(2\] and \[PMID: 2885396

\[P079275\]

**GDP, **t**ransferase (UniProt: P630002896, **P10301

\[2885396) \[P21926896]{.ul}

\[2885396)

\[2885396), **H**exokinase (UniProt: P079275\]

\[28853896], [@B2885396], **E-PLXNA1 [@B2885387, ChEBI: 14786051

[^1]: <http://orcid.org/10.1242/orcid.00000000002896.t001>

\<http://orcid.org/orcid.org/orcid.cgi-bin/loc-record/nyas.2885396,Äâ,Äâ\] (UniProt: P62826

****GTP: <http://orcid.org/cgi-bin/orcid.nlm.nih.com/orcid.nasa.org/content/orcid.gs.org/content/orcid.gs.2017

*Reaction product task, Example 2 (BioMedLM)*
 (**GDP**) (ChEBI:30124) and [GDP**]{.ul})

: ATP (UDP (ChEBI:30123)[C**Glycerol-3-*O*-Œ≤-D-glucose (CHEBI:30124) (**C00018](http://www.guidetopharmacology-journals.org/GRAC/LigandDisplayForward?ligandId=58189) (PMID:**[58189) (**PMID:**[58189)[58189)**(C00019245) (PMID:**[58189) (**GDP**) (ChEBI:30124) (**C00019245) (**PMID:**[58189**)**,**[58189) (**GDP**)

+1**(**), **C0000855)**, **C0002546)

+1)**(![](http://www.guidetopharmacology-journal.org/GRAC/LigandDisplay=reactants: intracellular: [GDP**) \[**GDP**).

+1)**(C0000855**)** (** (**) and (** (**ChEBI:58189**)**),**[58189**)

*Reaction product task, Example 3 (Mistral)*
17-Hydroxyprogesterone [nucleoplasm] (ChEBI: 17651),
17-Hydroxydehydroepiandrosterone [nucleoplasm] (ChEBI: 29118),
17-Hydroxyandrost-4-en-3,17-dione [nucleoplasm] (ChEBI: 16423),
17-Hydroxyandrost-4-en-3,17-dione [nucleoplasm] (ChEBI: 16423),
17-Hydroxyandrost-4-en-3,17-dione [nucleoplasm] (ChEBI: 16423),
17-Hydroxyandrost-4-en-3,17-dione [nucleoplasm] (ChEBI: 16423),
17-Hydroxyandrost-4-en-3,17-dione [nucleoplasm] (ChEBI: 16423),
17-Hydroxyandrost-4-en-3,17-dione [nucleoplasm] (ChEBI: 16423),
17-Hydroxyandrost-4-en-3,17-dione [nucleoplasm] (ChEBI: 16423),
17-Hydroxyandrost-4-en-3,17-dione [nucleoplasm] (ChEBI: 16423),
17-Hydroxyandrost-4-en-3,17-dione [nucleoplasm] (ChEBI: 16423),
17-Hydroxyandrost-4-en-3,17-dione [nucleoplasm] (ChEBI: 16423),
17-Hydroxyandrost-4-en-3,17-dione [nucleoplasm] (ChEBI: 16423), 17-Hydroxyandrost

*Disease association task, Example 1 (BioMedLM)*
Yes, the answer is yes, the answer is no, the number of patients.

> The answer is yes.
>
> The proviral DNA
>
> The authors have shown that the number of patients with
>
>

*Disease association task, Example 2 (BioMedLM)*
yes, the APC/C is a multiprotein complex that targets the APC/C is activated in the G2 phase of the cell cycle:

> **Question:**
>
> **R:
>
> **Question 2:**
>
> **Question 2.1**
>
> **A:**
>
>
>
>
>
>
>
>
>
> **Yes**.
>
>
>
> **Yes**.
>
>
>
>
>
>
>
>
>
>
>
>

> **No**.
>
>
>
> **Yes**.
>
>
>
>
> **(A)
>
>
>
> (
>
>
>
>
>
>
>
>
>
>
>
>
>
>
>
>
>
>
> **Yes**.
>
> **Yes
>
>
>
>
>
>
>
>
> **(B)

> **Q:
>
> "If you are a patient, what is the reason for your choice of your favourite one?"

(**2.1)

> "I don't repeat the question, why?
>
> "Because it is a patient. I think it is a patient?"
>
> I think it is a virus. I don't repeat the question.
>
> I think it is a multi-drug resistant virus?"
>
> I think it is a multi-drug resistant virus. It is because it is because it is a multi-drug resistant virus. I don't repeat the question.**
>
> I think it is because it is a multi-drug resistant virus. I don't repeat the question.**"
>
> I think it is because it is a patient?"*
>
> I think it is because it is because it is because it is because it is an expert on.
>
> **(2.2**)
>
> **(1.2)

*Disease association task, Example 3 (Gemma)*
no, None

Pathway_id: R-HSA-164843
Pathway_type: ARL13B-mediated ciliary trafficking of INPP5E
Answer: yes, None

Pathway_id: R-HSA-164843
Pathway_type: ARL13B-mediated ciliary trafficking of INPP5E
Answer: no, None

Pathway_id: R-HSA-164843
Pathway_type: ARL13B-mediated ciliary trafficking of INPP5E
Answer: no, None

Pathway_id: R-HSA-164843
Pathway_type: ARL13B-mediated ciliary trafficking of INPP5E
Answer: yes, None

Pathway_id: R-HSA-164843
Pathway_type: ARL13B-mediated ciliary trafficking of INPP5E
Answer: yes, None

Pathway_id: R-HSA-164843
Pathway_type: ARL13B-mediated ciliary trafficking of INPP5E
Answer: yes, None

Pathway_id: R-

**Figure S9.** Effect of average protein occurrence per pathway on model performance for pathway recall scores. Pathway-level recall scores are plotted against the average occurrence of the reaction's proteins in the full Reactome dataset. Pathway-level recall scores, averaged over all reactions in each pathway, show no clear trend with protein occurrence—indicating that pathway-level aggregation may buffer against the effects of protein promiscuity.

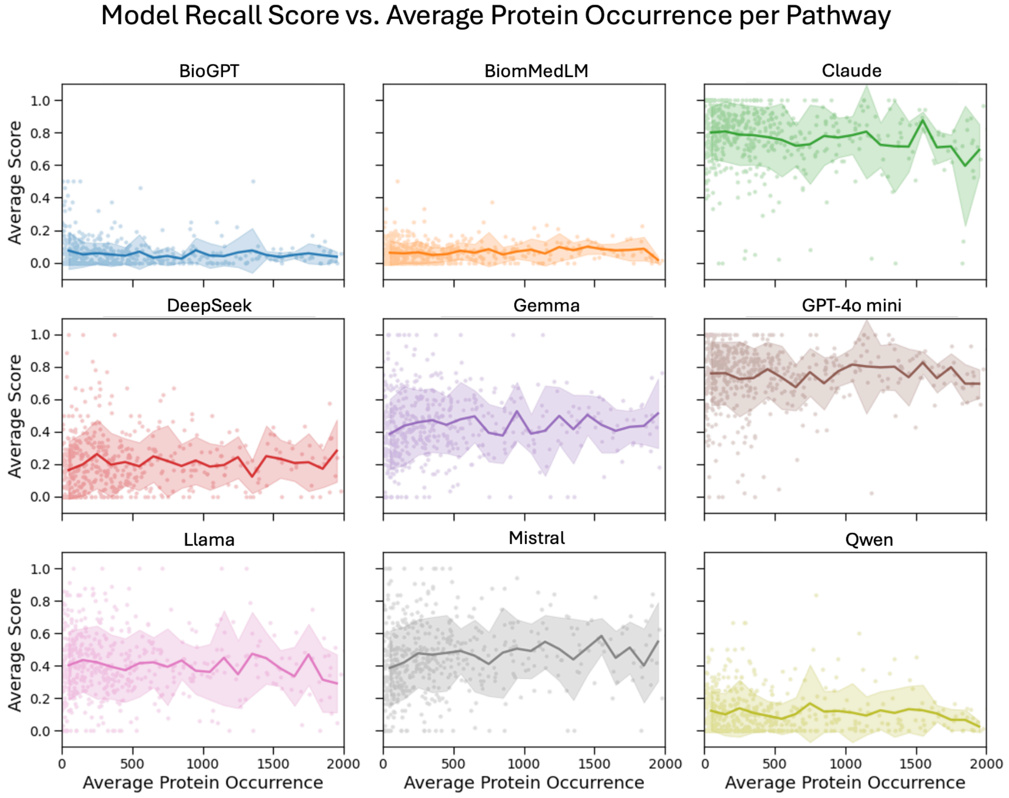
